# Non-Invasive Cancer Detection in Canine Urine through *C. elegans* Chemotaxis

**DOI:** 10.1101/2022.04.29.490074

**Authors:** Chan Namgong, Jong Hyuk Kim, Myon-Hee Lee, Daniel Midkiff

## Abstract

Cancer is the leading cause of death of companion animals, and successful early treatment has been a challenge in the veterinary field. We have developed the Non-Invasive Cancer Screening (N.C.S.) Study to perform cancer detection through the analysis of canine urine samples. The test makes use of the strong olfactory system of the nematode *C. elegans*, which was previously shown to positively respond to urine samples from human cancer patients. We performed a proof-of-concept study to optimize the detection capability in urine samples obtained from dogs with naturally occurring cancers. In this study, we established a scale for identifying the cancer risk based on the magnitude of the chemotaxis index of *C. elegans* towards a canine urine sample. Through validation, the N.C.S. Study achieved a sensitivity of 85%, showing that it is highly sensitive to indicating the presence of cancer across multiple types of common canine cancers. The test also showed a 93% specificity to cancer samples, indicating a low rate of over-identifying cancer risk. From these results, we have demonstrated the ability to perform low-cost, non-invasive cancer detection in companion animals, a method which can increase the ability to perform cancer diagnosis and treatment.

## 1 Introduction

There are over 200 million companion animals (dogs, cats, and horses) in the United States, and cancer is the leading cause of death among them (1). Approximately 1 in 4 dogs and 1 in 5 cats will develop cancer in their lifetimes, according to the Veterinary Cancer Society (2). Cancer in companion animals is difficult to treat successfully because few symptoms are evident in its early stages, and by the time symptoms become apparent, the cancer is usually advanced with a high mortality. It has been shown that approximately half of all canine cancers are treatable if diagnosed early enough, and new therapeutic approaches are continuously being established (3-6). Existing tests are available for cancer screening (7) but can be too expensive or invasive to be conducted regularly for some dog owners. Thus, there is an urgent need for the development of a novel economical and non-invasive method for cancer screening to increase the probability of successful treatment.

*C. elegans* is a simple multicellular organism that is often used as a model to study biological phenomena such as cellular signaling, neural development, and aging in higher multicellular animals (8). Breeding isogenic populations of *C. elegans* is straightforward, as nearly all animals in a wild-type population are hermaphrodites that reproduce through self-fertilization and are fed a diet of *E. coli. C. elegans* possesses a highly sensitive olfactory system to navigate its environment and detect food through the identification of chemical cues (9,10). *C. elegans* encodes at least 1,500 predicted G-protein-coupled receptors (GPCRs) (11). Some of these GPCRs are olfactory receptors that underlie the worm’s odor detection capabilities. *C. elegans* has an excellent sense of smell, and possesses approximately 1.5 times as many different types of olfactory receptors as a dog (12). Nematodes such as *C. elegans* rely on their strong sense of smell to search for food and navigate their environments (10). Once *C. elegans* detects an attractive odorant, it aligns with the chemical odorant and travels toward it (a process known as chemotaxis). This acute sense of smell allows for *C. elegans* to detect distinct volatile organic compound (VOC) profiles within animal urine.

The volatilome is the collection of VOCs which are present in the outputs of a biological organism (13). Cancer cells are known to emit VOCs that produce an odor that is distinguishable from that of non-cancer patients (14,15). Changes in the volatilome of specimen in animals affected with cancer have been measured using both gas chromatography and mass spectrometry (16,17). These odorant signatures are detectable in samples acquired from animals such as dogs (18) and mice (19), and thus could serve as a marker for identifying cancer. It has been shown that *C. elegans* can quantitatively detect the presence of signature VOCs in both in culture media of cancer cells *in vitro* and in urine samples of human cancer patients through chemotaxis assays and through calcium imaging of the AWC neuron (20–23). However, *C. elegans* has not yet been shown to identify cancerous VOC signatures in canine urine samples.

Here, we conducted the Non-Invasive Cancer Screening (N.C.S.) Study to measure the differences in *C. elegans* chemotaxis between urine samples from canine cancer patients and urine samples from healthy dogs with no diagnosed cancer. In the first part of the N.C.S. Study, we acquired initial data used to develop a screening method that identifies increased cancer risk through assays of canine urine samples. The study assesses multiple replicates of plate-based chemotaxis assays to measure the olfactory response through a mean chemotaxis index (CI). Based on these results, a risk assessment is made based on how the index relates to that of previously measured cancer and non-cancer samples. To validate the performance of our method, we assessed its ability to identify increased cancer risk using urine samples from dogs with four common types of canine cancer. In doing this, we demonstrate the potential for accurate, rapid, and non-invasive screening for cancer risk using urine samples from canine veterinary patients.

## 2 Materials and Methods

### 2.1 Canine Urine Samples

An initial set of canine urine samples was obtained from Triangle Veterinary Hospital (Durham, NC), Lake Pine Animal Hospital (Apex, NC), New Light Animal Hospital (Wake Forest, NC), Bull City Veterinary Hospital (Durham, NC), Knightdale Animal Hospital (Knightdale, NC), and from the Ohio State University Center for Clinical and Translational Science. Upon acquisition, urine samples were immediately stored at-20 □C until assays are conducted. Each specimen was aliquoted into 100 μL portions to minimize repeat freezing and thawing each time an assay is performed.

### 2.2 Maintenance of *C. elegans*

*Caenorhabditis elegans* strain N2 and *Escherichia coli* strain HB101 were obtained from the *Caenorhabditis* Genetics Center (University of Minnesota). *C. elegans* was age-synchronized by standard bleaching protocols and was cultured at 20 □C on nematode growth media (NGM) plates seeded with HB101 bacterial lawns. NGM plates were purchased pre-poured from LabExpress (Ann Arbor, MI).

### 2.3 Chemotaxis Assays

Assays are performed using well-fed age-synchronized populations of N2 worms grown at 20 □C for three days, and are conducted on CTX plates (2% Agar, 5 mM KPO4 buffer at pH 6, 1 mM CaCl_2_, 1 mM MgSO4) which were purchased pre-poured from LabExpress (Ann Arbor, MI). Urine samples were thawed and diluted at a ratio of 1:2 of urine to CTX buffer (5 mM KPO4 buffer at pH 6, 1 mM CaCl_2_, 1 mM MgSO_4_). Urine samples were centrifuged and were then mixed at a ratio of 2:1 diluted urine to 1 M sodium azide. The urine mixture was then spotted at the “plus” marks on each chemotaxis plate. Control buffer mixture was prepared using a 2:1 ratio of CTX buffer to 1 M sodium azide and was spotted at the “minus” marks on each chemotaxis plate (Supplementary Figure 1). Young adult worms were washed from NGM plates using M9 buffer into conical tubes and were allowed to settle. Worms were then washed three times with CTX buffer to remove traces of the bacterial food source. Approximately 75-100 worms were placed at the center of each plate, which were placed in a 23 □C incubator. After one hour, each plate was removed and placed on a backlight, and an image of each plate was acquired using an iPhone X digital camera.

### 2.4 Data Acquisition

Data was collected by manually counting the animals in each quadrant using Fiji ImageJ software. Replicates are discarded if one of the three conditions are met: (1) if the total for all four quadrants is less than 55 (2); if the highest total quadrant exceeds the sum of the remaining three quadrants; (3) if the quadrant across from the highest total quadrant has fewer than half the animals of any other quadrant (Supplementary Figure 2). Then, the CI is calculated using the following formula, where Qn is the number of worms in the nth quadrant:

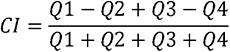

A mean CI is calculated from the replicates for each assay that are not discarded.

#### 2.5 Statistical Analysis

Differences between cancer risk groups were assessed using the Welch t-test for data sets with unequal variance. Thresholds for cancer risk assessment were drawn to optimize the accuracy of the data analysis. 95% confidence intervals were specified based on the following formula

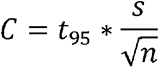

Where C is the confidence interval, t_95_ is the t-score, s is the sample standard deviation, and n is the sample size.

## 3 Results

### 3.1 Developing The A.C.D. Test to Distinguish Cancer and Non-cancer Urine Samples

We developed the N.C.S. Study based on chemotaxis data that was previously generated showing a slight preference of *C. elegans* for urine samples acquired from cancer patients, as opposed to a slight aversive response to urine from non-cancer patients (20–22). Our first approach was to determine if the procedure previously used for human cancer detection could be applied to canine urine. While performing chemotaxis, we found that individual replicate outliers could cause drastic swings in the calculated mean CI. For this reason, replicates that deviated strongly from the other replicates in the assay were discarded. We chose to discard CI with a difference greater than 0.25 from the closest other replicate within the assay to reduce the distortion from extreme data points on the mean chemotaxis value. Plates with fewer than 55 worms in the four quadrant boundaries were discarded, as plates with fewer worms tended to yield a wider range of CI values, leading to greater distortions in the mean. We also discarded replicates with unusual distributions (Supplementary Figure 2). We defined plates as yielding outlier results when the total number of worms in one quadrant exceeded the total number in all three other quadrants combined, or when the number of worms across from the highest total quadrant is less than 50% the total of any other quadrant. These arrangements indicated migration either towards or away from a particular quadrant rather than a particular chemical stimulus and did not provide reliable data for calculating a mean CI.

Through these optimizations, we developed the N.C.S. Study to accurately identify urine samples from canine cancer patients (Figure 1). Previous chemotaxis assays have used anywhere from three to six replicates to determine the mean CI (20–22). For canine urine assays, we often found variance in the response and magnitude in individual replicates within an assay (Supplementary Table 1). We found that it was necessary to acquire at least five non-discarded replicates for one urine test, four replicates if all are positive or negative. From the CI replicates which were not discarded, we calculated the mean CI, which was then used to assess the level of risk.

**Figure 1:**
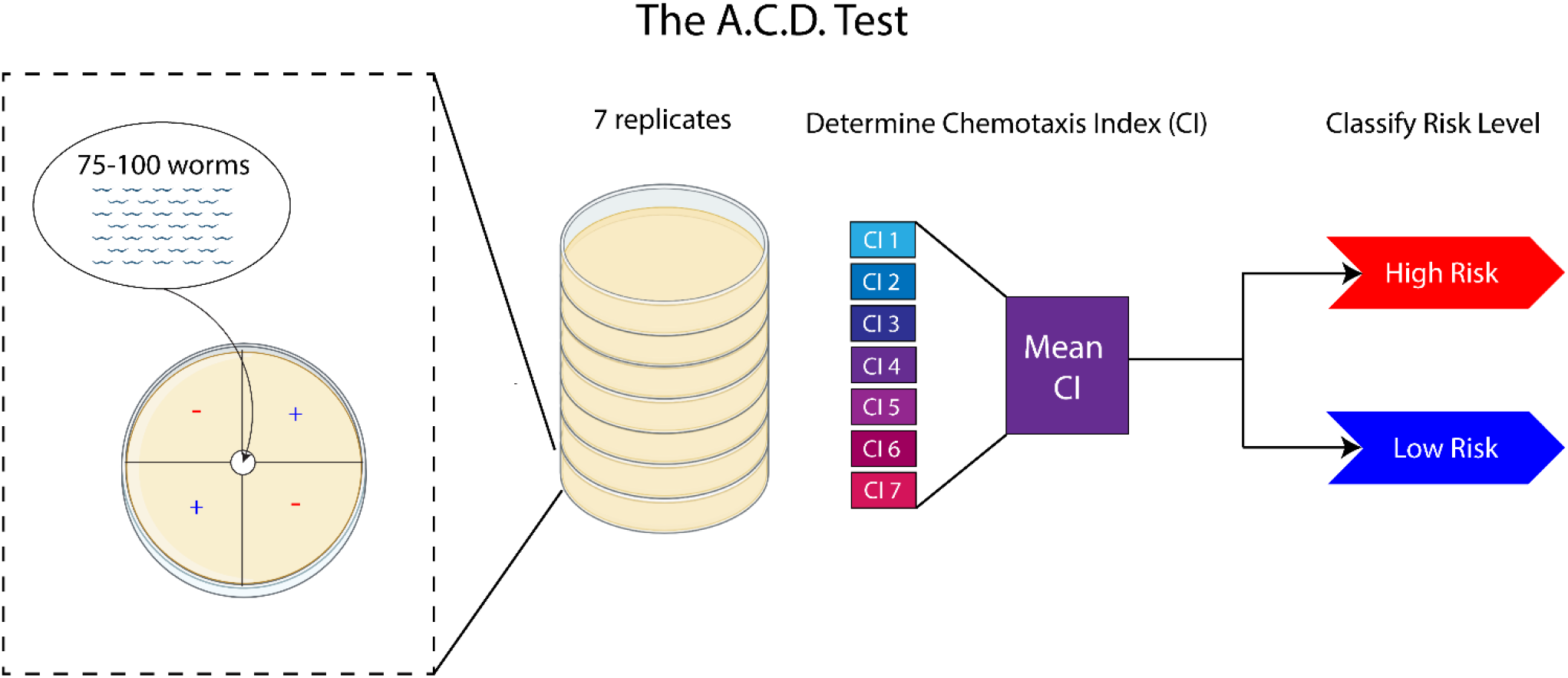
The A.C.D. Test is conducted by placing 75-100 worms on each assay plate. A total of seven assay replicates are conducted, from which a mean CI is calculated and used to assess the level of cancer risk.

### 3.2 Determining the Level of Cancer Risk for the A.C.D. Test

We performed tests on a series of cancer and non-cancer urine samples to determine if cancer can be detected through positive chemotaxis towards canine urine samples. We initially performed assays on a total of eight cancer samples and fourteen non-cancer samples. We found that *C. elegans* was much more strongly attracted to cancerous urine samples than non-cancer samples (Figure 2A). From these results, we set a threshold for elevated cancer risk determined from a one-way Student t-distribution of the tested non-cancer samples (α=0.005). Mean CI values less than or equal to 0.038 are classified as “low risk”, as that is the range of about 85% of non-cancer samples, while we designated results above the threshold as “moderate to high cancer risk.” Through this approach, we achieved an 88% sensitivity for cancer detection, and a 93% specificity for correctly classifying non-cancer samples (Figure 2B, Supplementary Table 2). We also ran replicates of a cancer and noncancer sample which indicated replicable outcomes of the assay risk classification (Supplementary Table 3). Based on these results, we have shown a preliminary ability to classify the cancer risk of canines through *C. elegans* chemotaxis.

**Figure 2:**
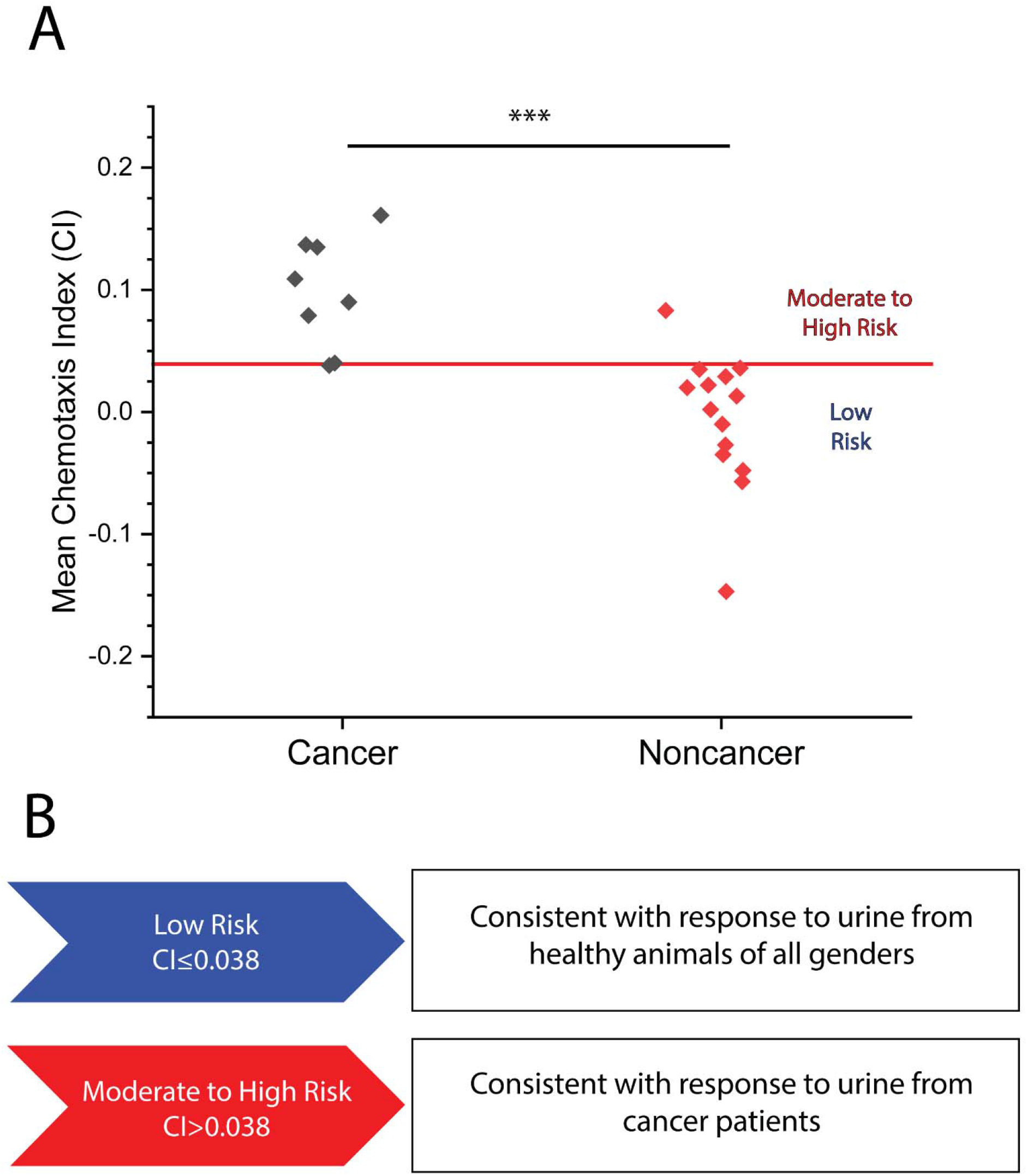
(A) Mean CI plotted for eight cancer and 14 non-cancer samples for which the A.C.D. Test was conducted. A mean CI of 0.099±0.038 for cancer samples versus a mean CI of-0.006±0.032 in non-cancer samples (p=0.0002) Red line indicates moderate to high cancer risk classification threshold. ***p<0.001 (B) Levels of cancer risk set at the following range: Low Risk (<0.038) and Moderate to High Risk (>0.038)

### 3.3 Assessing Detection Rate of Four Common Canine Cancers

To further determine the accuracy of the NC.S. Study at detecting the presence of cancer, we performed assays on ten samples of each of four different types of cancer that are commonly diagnosed in domestic dogs: lymphoma, mast cell tumor, melanoma, and hemangiosarcoma (24) (Figure 3, Supplementary Tables 4 and 5). We found that all samples yielded a higher mean CI than for non-cancer samples. By combining the data acquired from these forty samples with that in the preliminary data set, we found that the N.C.S. Study yielded a sensitivity of 85% of identifying at least a moderate risk of cancer in each confirmed cancer patient, as compared to the 7% of non-cancer samples identified as at least a moderate cancer risk (Table 1). Overall, by combining all measured CI values for cancer and non-cancer samples, we achieved an accuracy of 87%. Our results also showed a statistically significant difference between the mean CI for each type of cancer and the non-cancer samples.

**Figure 3:**
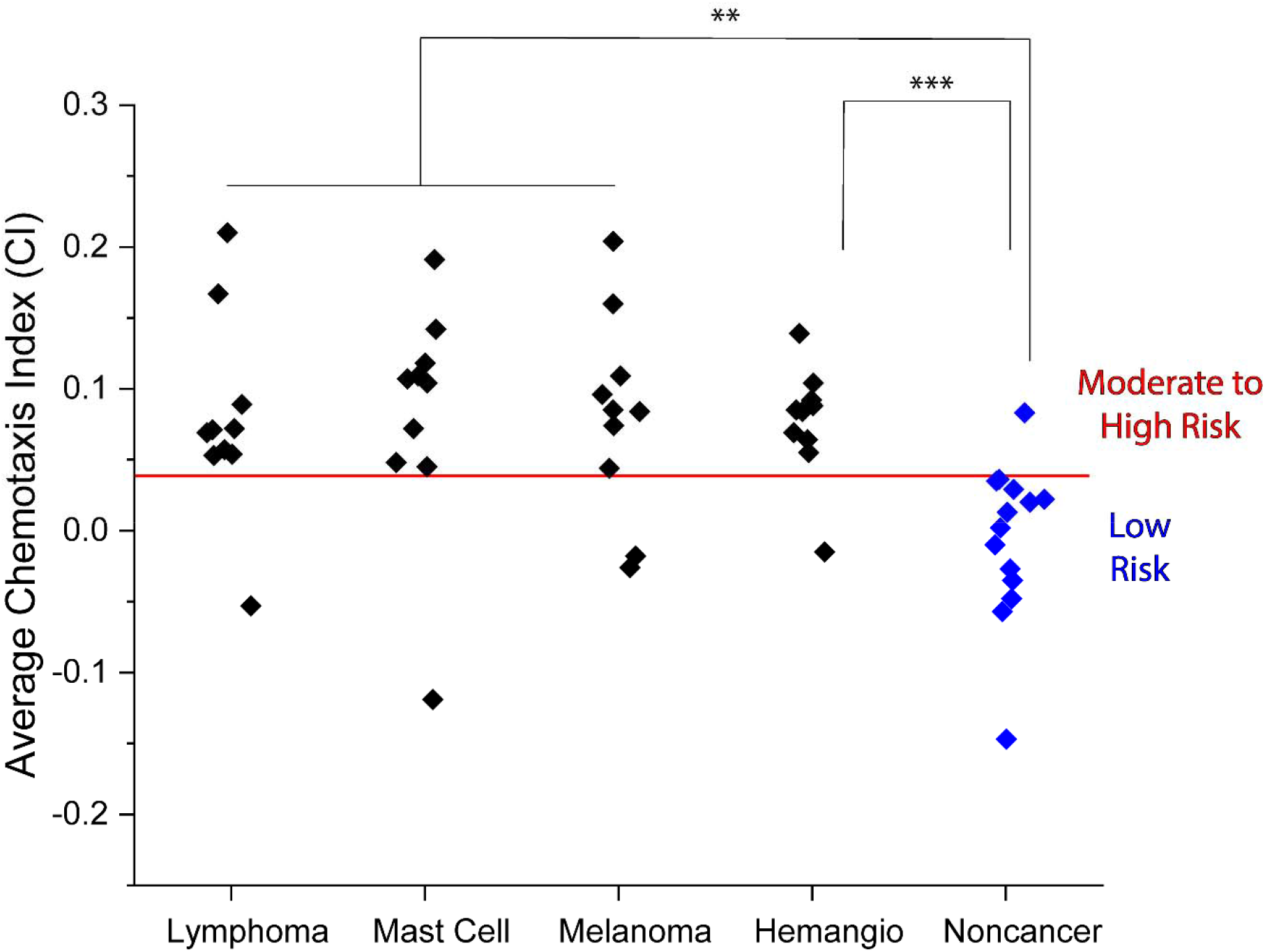
Average CI for urine samples obtained from lymphoma, mast cell tumor, melanoma, and hemangiosarcoma patients as compared to non-cancer samples. Lymphoma: 0.079±0.050, mast cell tumor: 0.081±0.059, melanoma: 0.081±0.051, and hemangiosarcoma: 0.077±0.028 versus non-cancer:-0.006±0.032). Red line indicates moderate to high cancer risk classification threshold. **p<0.01, ***p<0.001

**Table 1:**
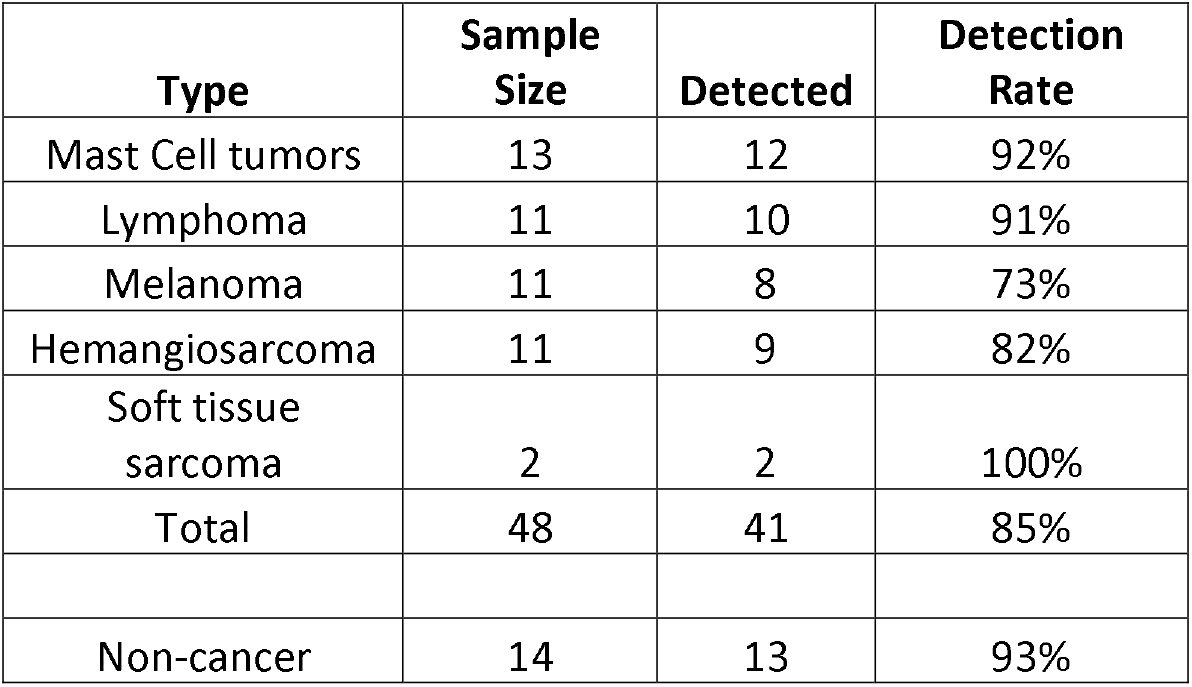
Data summary for each classification of cancer versus non-cancer urine samples.

## 4 Discussion

In this study, we assessed how the presence of cancer in canines affects the odor of urine samples as perceived by the nematode *C. elegans* and to determine how it compares to the detected odor in human cancer and non-cancer patients. Additionally, while we were able to replicate the significantly more attractive chemotaxis response in cancer-positive canine samples, we observed a mean CI across all non-cancer samples that was approximately zero. Urine is composed of a mix of salts, minerals, hormones, and other chemicals, all of which contribute to the chemotaxis response for a sample (25,26). Thus, the difference in composition between human and canine urine could yield differing mean chemotaxis responses for non-cancer samples.

Each of the four types of cancer for which we screened ten samples was detected at a high rate in the N.C.S. Study, with no cancer type having a sensitivity below 70%. We thus observe that risk of multiple types of cancer can be identified through urine chemotaxis assays. We also performed two assays on samples from patients with soft-tissue sarcoma, a type of cancer which is less prevalent but still occurs in some canines. We found that both samples were detected by chemotaxis assays, showing that it is another type of cancer that potentially can be detected at a high rate using urine chemotaxis assays. While the test can identify the presence of cancer at a high rate, the test does not give any information on which type of cancer a patient has developed. Moreover, a positive result in our test does not constitute a cancer diagnosis, but a warning of a risk that cancer may be present at a certain stage in a canine patient. Further diagnosis and monitoring of symptoms is necessary to confirm the presence of cancer and its type. In addition, it has previously been shown that attractive VOCs are present in the urine during both early and late stages of cancer(20,22), so it is likely that in future assays, we can demonstrate the ability to screen for cancer at the earliest stages.

The N.C.S. Study yielded performance metrics that fell into the range of accuracies that was previously determined for human cancer samples. The performance metrics were slightly weaker than what was previously measured by Hirotsu et al, who detected several types of cancer from human urine using *C. elegans* chemotaxis with a sensitivity of 96% and a specificity of 95%, yielding an overall accuracy of 95% (20). However, the performance was comparable to that achieved by Lanza et al. through chemotaxis (accuracy of 86%) (21) and exceeded that achieved by Thompson et al (accuracy of 70%) (22). Thus, our results lend evidence to the hypothesis that patterns in cancerous VOC signatures that have been well-studied in humans are comparable with that in canines. Since VOCs have also been detected in urine samples from cancerous mice (23), there is strong evidence that similar methods can be utilized to detect cancer in feline, equine, and other mammalian veterinary patients. However, further studies are necessary to determine for which mammalian species this test can be applied, and how effective this method is at distinguishing cancer samples from non-cancer samples.

While we were able to measure a significant difference in chemotaxis towards cancer and non-cancer urine samples, in some instances we found a high level of variance between individual replicates within an assay. While we have not identified the source of this variation, these results indicate that the locomotion of *C. elegans* is highly random. However, we also observed that cancer samples producing a strongly positive chemotaxis index will have few if any negative replicates. Nonetheless, assays with a wide range of replicate CI values are more likely to produce different results when repeated. For this reason, by achieving a stronger chemosensory attraction towards cancer urine samples, or by reducing the variance of individual replicates, the accuracy and replicability of cancer detection through odor can be improved. It has previously been shown that a response to positive volatile odorants present in cancerous urine samples can be identified through calcium gradients in the *C. elegans* AWC sensory neuron (27). This characteristic calcium gradient is indicative of the presence of an attractive odorant and is strongly distinguishable from the low gradient that has been measured for noncancer samples. Moreover, a much lower noise level is achieved using this method as compared to odor detection through chemotaxis. We hypothesize that by measuring calcium gradients, we could achieve an even higher accuracy of cancer detection from canine urine samples.

In recent years, the number of pet owners in the United States and around the world has undergone a steady increase. This, combined with the strong emotional bonds that owners have with their pets, creates a higher demand than ever for treatments to protect pets from life-threatening illnesses. Collection of urine samples is routine and non-invasive, and the test can be conducted accurately at a high rate in a basic laboratory setting. By detecting more attractive urine chemotaxis in multiple common cancer types, we present evidence towards urine odor as an effective method for cancer screening in canines.

## 5 Conflict of Interest

Authors Chan Namgong and Daniel Midkiff were employed by the company Animal Cancer Dx.

The authors declare that this study received funding from Animal Cancer Dx. Employees of Animal Cancer Dx had the following involvement with the study: developed the research idea, performed and conducted experiments, and wrote and edited the article.

## 6 Author Contributions

Daniel Midkiff grew and maintained stocks of *C. elegans* and conducted all chemotaxis experiments and was the primary author of the article. Chan Namgong developed the research idea, assisted in experimental development, and collected urine samples. Myon-Hee Lee assisted in experimental design and setup and contributed to the writing and editing of the article. Jong Hyuk Kim contributed to data interpretation and the writing and editing of the article.

## 7 Funding

This work was supported by Animal Cancer Dx to C.N. and D.M., and the National Science Foundation (IOS-2132286) to M.H.L.

### 8 Acknowledgments

This publication was supported, in part, by the National Center for Advancing Translational Sciences of the National Institutes of Health under Grant Number UL1TR002733. The content is solely the responsibility of the authors and does not necessarily represent the official views of the National Institutes of Health.

Strains were provided by the CGC, which is funded by NIH Office of Research Infrastructure Programs (P40 OD010440).

## 9 Contribution to the Field Statement

Cancer is one of the leading causes of death in companion animals and occurs in approximately one quarter of all domestic dogs at some point in their lives. Despite its prevalence, it has been an immense challenge to treat cancer in canines, as it frequently diagnosed after symptoms are apparent. At this advanced stage, the cancer has often spread too far for successful treatment. In this publication, we have demonstrated that cancer in canines can be detected by identifying the presence of volatile compounds which yield a distinguishable odor. This odorant signature can be identified through the response of organisms such as the nematode *Caenorhabditis elegans*. Here, by measuring the positive response of *C. elegans* to volatile odorants in canine urine samples, we were able to assess cancer risk. These results are replicable in four types of cancer that are common in companion dogs. Through these results, we show that odor detection assays can serve as an accurate, rapid, and noninvasive cancer screening method in dogs. By performing more frequent cancer screening throughout a dog’s life, cancer can be detected at an earlier stage and increase the odds of successful treatment.

## Supplementary Material

### 1 Supplementary Figures

**Supplementary Figure 1.**
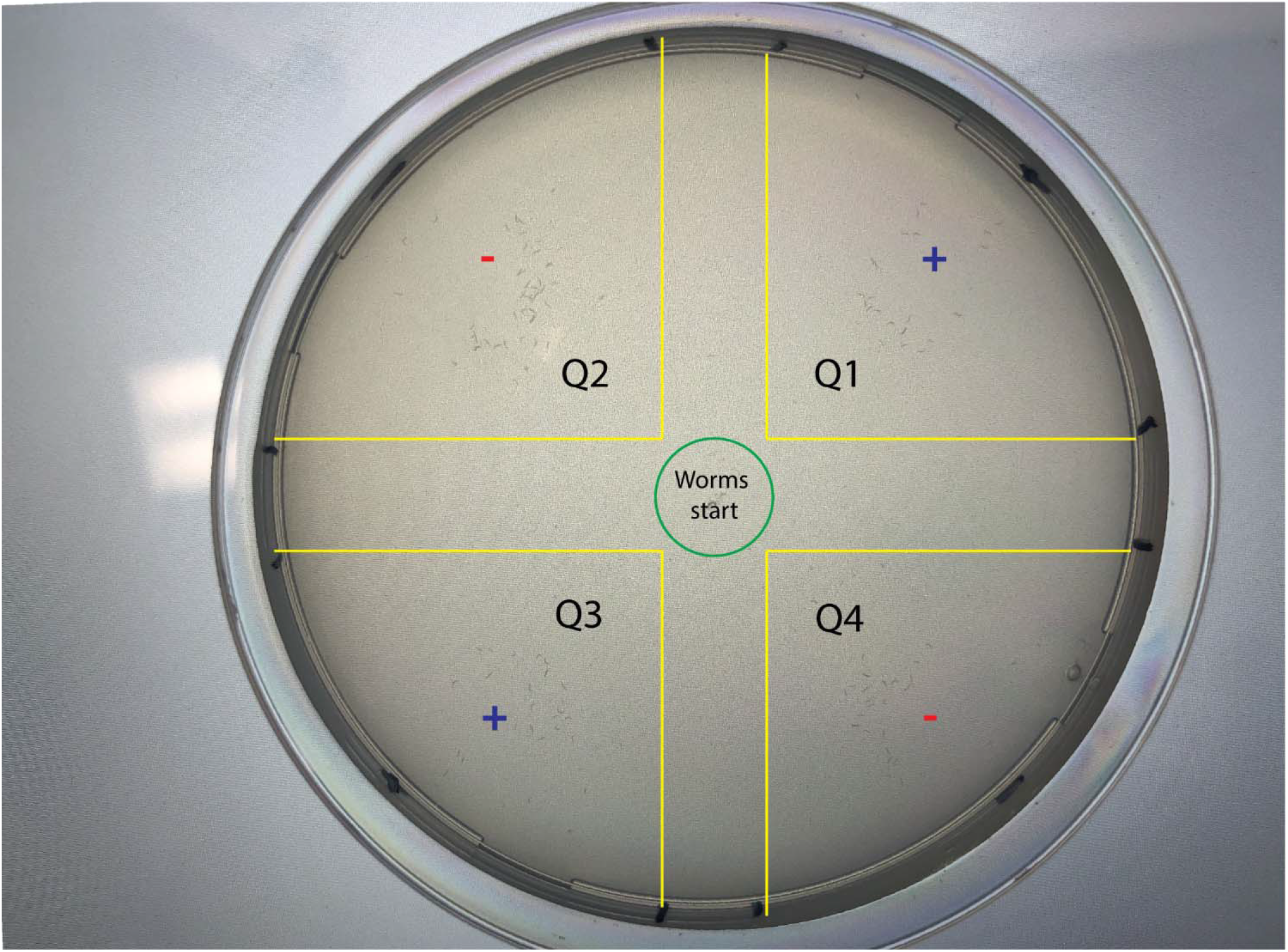
Each assay plate is divided into four quadrants. The positive quadrants (Q1, Q3) are marked with a “plus” sign, which designates the location of the tested urine sample. The negative quadrants are marked with a “minus” sign, which designates the location of the control buffer. *C. elegans* are placed at the center of the green circle at the beginning of the assay. After the assay is complete, animals are counted in each quadrant within the bounds of the yellow lines to avoid counting those that are close to the quadrant borders or that remain at the center of the plate.

**Supplementary Figure 2.**
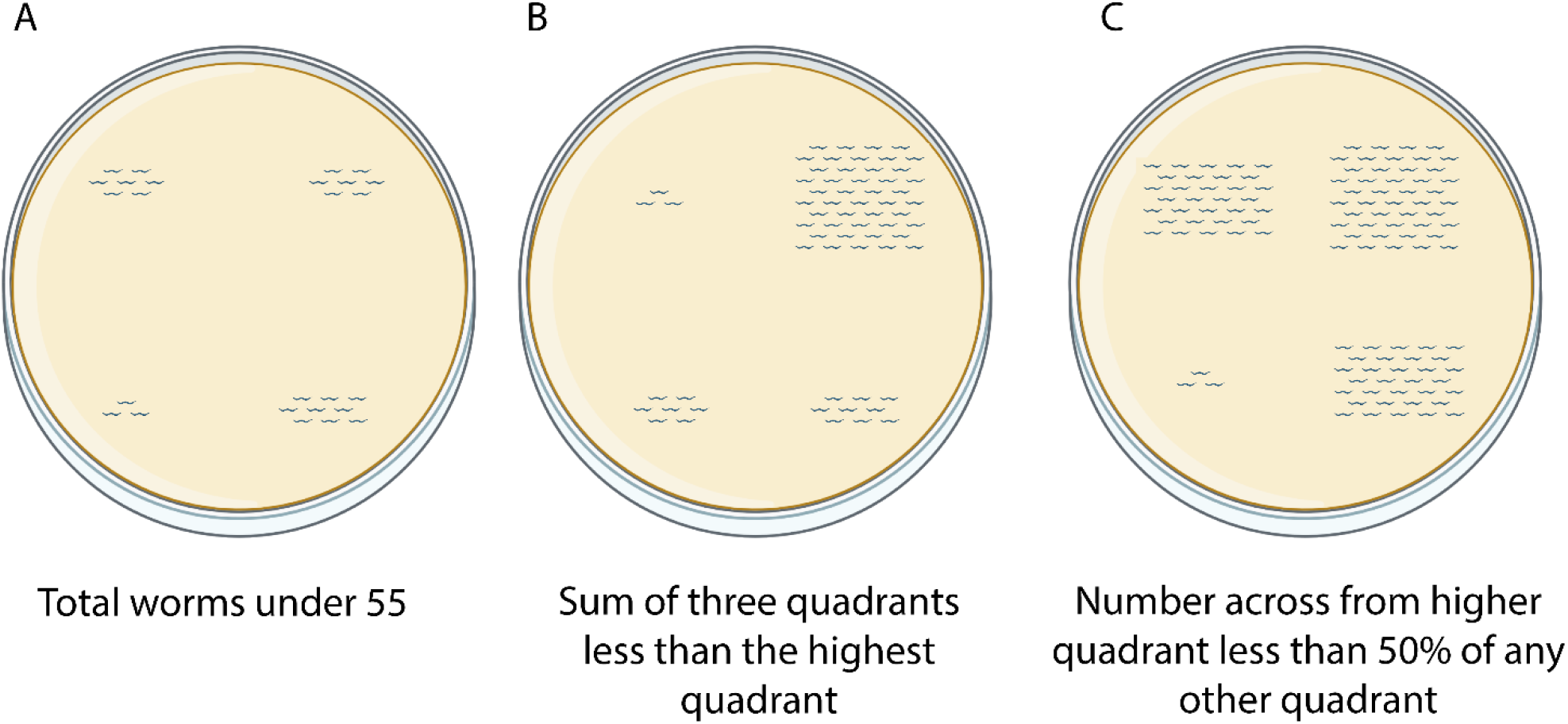
Examples of conditions in which plates are discarded. (A) The total number of animals on the plate is less than 55:

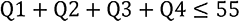 (B) The sum of the total animals in three quadrants is less than the total number of animals in the fourth quadrant, e.g.

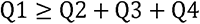 © The number of animals across from the quadrant with the highest total has 50% or less than the number of animals on any other quadrant, e.g.

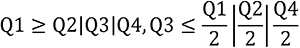

### 2 Supplementary Tables

**Supplementary Table 1.** CI replicates for initial set of cancer and non-cancer assays. The CI for each non-discarded replicate is listed beneath the name of each cancer and non-cancer patient.

**Supplementary Table 2.** Chemotaxis data summary for initial cancer and non-cancer data comparison.

**Supplementary Table 3.** Sample assay repeats for dog patients with (P03) and without (P09) diagnosed cancer. Each of the assay repeats shows a mean CI above and below the moderate risk threshold for cancer and non-cancer patients, respectively.

|  | P03 (Mast Cell Tumor) |  |  | P09 (Non-cancer) |  |  |
| --- | --- | --- | --- | --- | --- | --- |
|  | Assay 1 | Assay 2 | Assay 3 | Assay 1 | Assay 2 | Assay 3 |
| Replicate 1 | 0.123 | 0.223 | 0.109 | -0.159 | -0.051 | 0.294 |
| Replicate 2 | 0.233 | 0.278 | 0.052 | -0.153 | 0.163 | -0.294 |
| Replicate 3 | 0.12 | -0.042 | 0.045 | -0.167 | -0.136 | -0.127 |
| Replicate 4 | 0.174 | 0.286 | 0.071 | 0.031 | 0.164 | 0.147 |
| Replicate 5 | 0.102 | 0.147 | 0.211 | 0.096 | -0.31 | -0.143 |
| Replicate 6 | 0.082 |  | 0.027 | 0.056 |  |  |
| Replicate 7 | 0.109 |  |  | 0.053 |  |  |
| Mean CI | 0.135 | 0.178 | 0.086 | -0.035 | -0.034 | -0.025 |

**Supplementary Table 4.** CI replicates for ten samples of lymphoma, mast cell tumor, melanoma, and hemangiosarcoma. The CI for each non-discarded replicate is listed beneath the name of each cancer and non-cancer patient.

**Supplementary Table 5.** Chemotaxis data summary for ten samples of lymphoma, mast cell tumor, melanoma, and hemangiosarcoma.

